# Intranasal photobiomodulation: an energy efficient paradigm for cortical and subcortical stimulation

**DOI:** 10.64898/2026.03.03.709361

**Authors:** Hannah Van Lankveld, Joanna X. Chen, Xiaole Z. Zhong, J. Jean Chen

## Abstract

Previous studies have shown the hemodynamic response to transcranial photobiomodulation (tPBM) in localized cortical regions during and after forehead irradiation. However, it is unclear if tPBM can reach deeper regions such as subcortical tissue. It is also unclear whether the manner of regional neurovascular coupling predominantly studied using tPBM extends to all brain regions. As an alternative to forehead delivery, intranasal PBM (iPBM) uses the pathway of the cribriform plate, which is thin and directly leads to the orbitofrontal cortex, rather than the prefrontal cortex in the case of forehead PBM. Thus, it is possible that iPBM can stimulate the brain more efficiently (i.e. with less power). In this study, healthy young adults underwent different iPBM protocols differing in wavelength, frequency and irradiance. We utilized functional magnetic resonance imaging (fMRI) to quantify regional blood oxygenation (BOLD) and perfusion. We further model the neurovascular interactions underlying the fMRI response. We uncovered three distinct temporal signatures, varying by brain region. Specifically, a significant response in the thalamus was observed, with a time-locked BOLD response. Overall, iPBM was found to be associated with much higher efficiency at eliciting BOLD fMRI responses than its forehead (tPBM) counterpart. Lastly, in addition to the expected dose dependence, there were extensive sex differences in the fMRI response to iPBM, surpassing those observed for tPBM. Collectively, these findings highlight the feasibility and efficacy of iPBM and establish a foundation for personalizing PBM protocols for optimal outcomes.

## Introduction

Photobiomodulation (PBM) (Mester et al. 1968) relies on the ability of light in the NIR range, (wavelengths of 630 nm - 1060 nm) to penetrate biological tissues (Tedford et al. 2015) and influence mitochondrial function (Karu et al. 2007), cerebral metabolism as well as neurovascular coupling (Tian et al. 2016). The absorption of NIR light by cytochrome-c-oxidase (CCO) leads to enhanced mitochondrial respiration, ATP production, nitric oxide dissociation, downstream vasodilation and neurovascular modulation (Thunshelle and Hamblin 2016), (Collman et al. 2008). In addition, PBM is known to modulate and activate ion channels, including calcium, potassium, sodium and transient receptor potential (TRP) channels, which further influence cellular metabolism, vascular responses and neural signaling (Zhang et al. 2024). The most common form of light delivery is via the forehead or superior skull (Salehpour et al. 2018). Our recent studies characterized the dose dependence of forehead tPBM in terms of not only laser parameters, but also of sex and skin tone (Van Lankveld, Mai, et al. 2025; Mathew et al. 2025). Our previous work has shown that tPBM elicits region-specific hemodynamic responses that extend beyond the stimulation site. Distinct temporal profiles were observed across regions of interest (ROIs), with some responses tightly following the stimulation period and others remaining elevated afterward. The study also demonstrated clear dose dependence across stimulation parameters and biological factors including skin pigmentation and sex. Importantly, using a combined functional magnetic resonance imaging (fMRI) measurement of blood blood-oxygenation-level-dependent (BOLD) imaging and cerebral blood flow (CBF), it was noted that certain ROIs exhibited changes in the relationship between these two signals over time, representing multiple possible neurovascular response scenarios, suggesting that tPBM can engage multiple metabolic and vascular pathways within a specific region. However, our work showed no response in subcortical areas. Although tPBM has demonstrated measurable physiological effects in cortical tissues, it remains uncertain whether sufficient light energy applied to the forehead can penetrate deeper brain regions, such as subcortical structures, as most human tPBM studies have focused on cortical regions. Moreover, these results are often interpreted within the framework of neurovascular coupling, but it is also unclear whether conventional neurovascular coupling extends to tPBM. Additionally, given that a small fraction of the NIR light penetrates the skull, it is of interest to find if an alternate delivery route sustains less loss.

In intranasal PBM (iPBM), light is delivered via the nasal cavity (Jiao et al. 2005), where thin bones and rich vascular networks may facilitate greater transmission of photons. iPBM was first reported in 1998 with early clinical applications emerging in the 2000s, and its feasibility has been demonstrated through Monte Carlo simulations (Cassano et al. 2019; Van Lankveld, Mai, et al. 2025). Moreover, early evidence from animal studies and clinical work suggests potential benefits for cognition, mood and neurodegenerative disorders (Chao 2019), (Salehpour et al. 2020). Recent literature has examined the cognitive response to iPBM but only in conjunction with tPBM in potential concussion and Alzheimer’s disease (Johnson et al. 2024), (Liebel et al. 2025)(Chao 2019)(Zomorrodi et al. 2019). Due to iPBM’s light path via the orbitofrontal cortex, it was suggested that iPBM can reach deep brain structures that were deemed unreachable using forehead stimulation. Moreover, due to the thinness of the cribriform plate compared to the skull, it is possible that the iPBM is more energy efficient than tPBM (Saltmarche et al. 2017) (n.d.). However, the effect of iPBM independently of tPBM has never been studied. Thus, there is a clear gap in the literature that compares tPBM and iPBM, and an even more stark scarcity in our understanding of the dose dependence of the brain’s response to iPBM. Dissecting the effect of iPBM from tPBM is important, along with understanding the effects of stimulation parameters. At the moment, all the aforementioned combinatorial studies of iPBM with tPBM applied the same wavelength (810 nm), and irradiance (25 mW/cm²) to iPBM. However, to date, there have been no published fMRI studies investigating signal responses to independent iPBM. Moreover, examining the interplay between the BOLD and CBF signals can reveal neurovascular mechanisms of the iPBM response (Van Lankveld, Chen, et al. 2025).

In this work, we address these gaps by directly measuring iPBM-evoked BOLD and CBF responses, assessing the spatial distribution and temporal signal characteristics both independently and then comparatively to our previous tPBM work. This study is the first fMRI assessment of iPBM in humans, evaluating the neurovascular coupling to infer underlying vascular and metabolic mechanisms. We also investigate the depth of stimulation across not only cortical regions, but also subcortical structures, which have been traditionally a challenge for brain stimulation (Liu et al. 2021; Herath 2026). Moreover, motivated by our simulation study (Van Lankveld, Mai, et al. 2025), we systematically assess dose dependence across multiple stimulation parameters (wavelength, irradiance and frequency) and sex. Structural and vascular differences between males and females suggest that sex could significantly modulate the distribution of the iPBM response. This work uses in vivo imaging to assess the value and accuracy of simulation-based predictions and advances our fundamental physiological understanding of PBM in the living brain.

## Methods

### Participants

We recruited 45 healthy young adults (age 20-32, 23 M/22 F) who were screened prior to their participation to ensure there was no history of neurological or physiological disorders, malignant disease, or the use of medications that could have influenced the study. The Baycrest Research Ethics Board (REB) approved the study, and all experiments were conducted in accordance with REB guidelines, with each participant providing written informed consent.

### PBM Instrumentation

The MRI-compatible lasers were provided by Vielight Inc., and the light was delivered through a 10-meter, 400μm optic cable to the MRI, while the laser system remained outside the scanner room. The light parameters, including wavelength, irradiance and pulsation frequency, were controlled from the console to turn the lasers on and off while the participant was inside the scanner. The participant remained blinded to the stimulation paradigm, the light was secured through a nose applicator, inserted inside the nasal cavity such that it was not visible to the participants, to avoid any placebo effects.

### MRI Acquisition

fMRI data was acquired using a Siemens Prisma 3 Tesla System (Siemens, Erlangen, Germany) with a 20-channel radio-frequency coil. For each participant, the following data was collected: (a) A T1-weighted structural image (sagittal, 234 slices, 0.7 mm isotropic resolution, TE = 2.9 ms, TR = 2240 ms, TI = 1130 ms, flip angle = 10°). (b) A dual-echo (DE) pseudo-continuous arterial spin labeling (pCASL) scan (Danny J. J. Wang, University of Southern California) to measure cerebral blood flow (CBF) and BOLD (TR = 4.5 s, TE1 = 9.8 ms, TE2 = 30 ms, post-labeling delay = 1.5 s, labeling duration = 1.5 s, flip angle = 90°, 3.5 mm isotropic resolution, 35 slices, 25% slice gap, total scan time = 12 minutes 15 seconds). Additionally, an M0 calibration scan was acquired with a TR of 10 s and a scan time of 45 s, while all other parameters remained unchanged.

Participants were scanned while viewing naturalistic stimuli to minimize variations in brain state across participants, thereby improving the reproducibility of the stimulation outcome (Gal et al., 2022).

### Stimulation Protocol

The stimulation parameters refer to the various settings of light that could influence the treatment’s effectiveness and safety. The study varied three irradiances (5 mW/cm², 7 mW/cm², and 9 mW/cm²), two wavelengths (808 nm and 1064 nm), and two pulsation frequencies (10 Hz and 40 Hz) across all participants to thoroughly evaluate the effects of iPBM on brain activity. The iPBM laser was carefully positioned within the right nostril and attached with a clip to keep it in place. This positioning targets the limbic system as the light passes through the cribriform plate and in the frontal sinus into the neural tissue. Each participant completed four separate iPBM-fMRI scans, each with a unique combination of stimulation parameters. Participants were assigned to one of three protocols, with each protocol comprising four scans. Out of the 45 participants, 15 were assigned to each protocol.

For each 12-minute scan, the stimulation paradigm was 4-min-OFF, 4-min-ON, 4-min-OFF, as shown in **Figure 1**. Each subject underwent 4 iPBM sessions each with a unique combination of dose parameters, as seen in **Table 1**. A period of at least 10 minutes was provided as a break before each subsequent iPBM session. In combination with the 4 minutes of pre-stimulus recording and 4 minutes of post-stimulus recording there was a minimum time of 18 minutes between stimulations.

**Figure 1:**
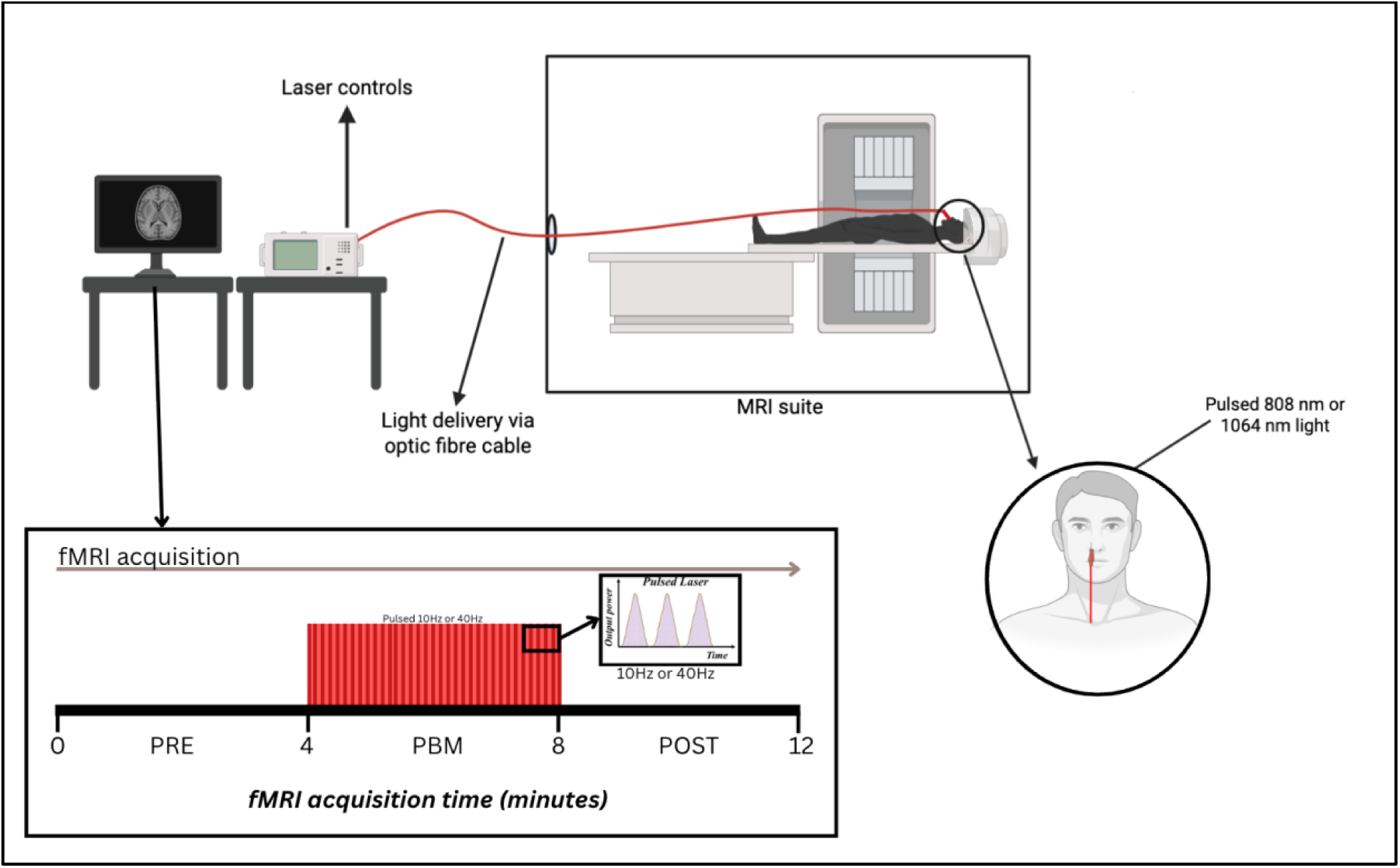
iPBM-fMRI setup; optic fibre cable carries the NIR light from the laser control system through a small hole into the MRI suite. The optic fibre is secured to the participants chest strap, fed through the RF coil and positioned on the right side of the forehead. From outside the MRI suite, the console allows the laser to be turned ON/OFF throughout the fMRI recording. fMRI acquisition follows a 4min-OFF, 4-min-ON, 4-min-OFF paradigm with pulsed NIR light at 10 Hz or 40 Hz.

**Table 1:**
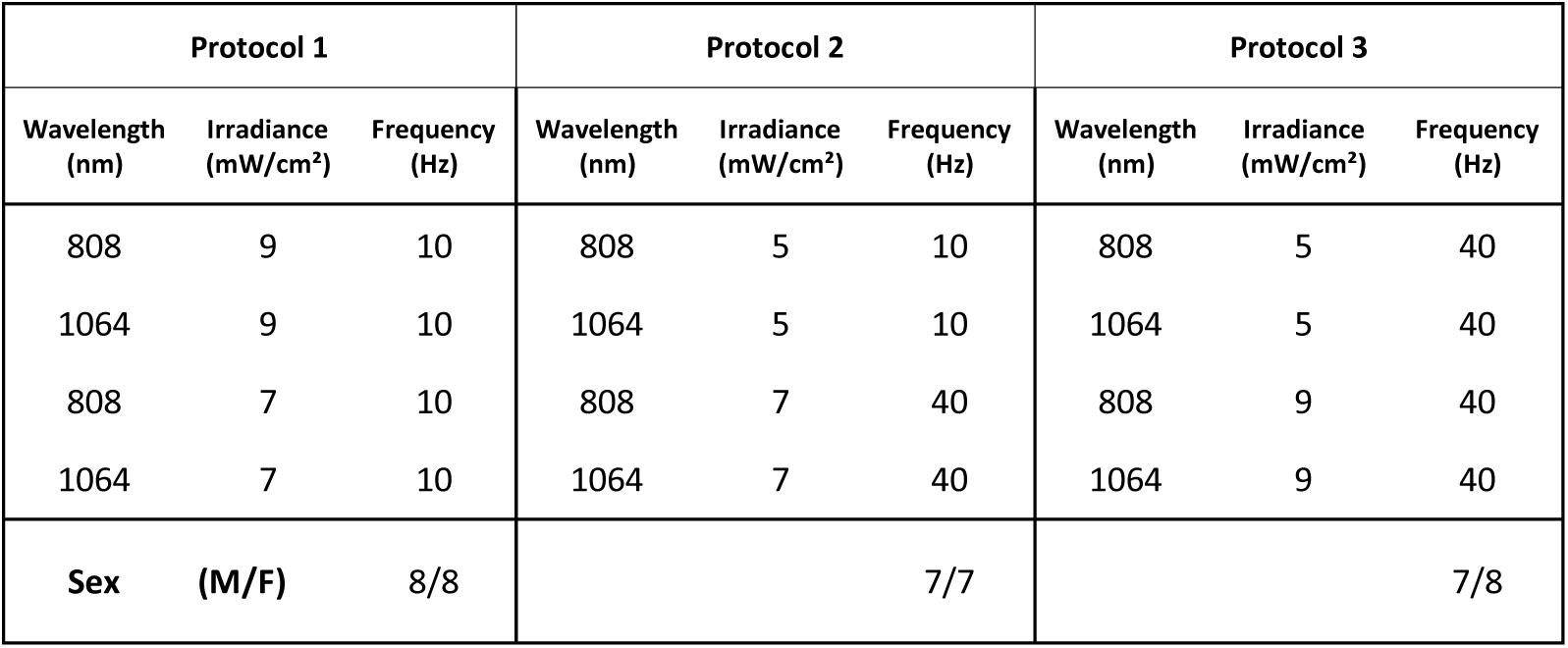
Stimulation paradigm for each protocol grouping.

### Data Preprocessing

BOLD-fMRI preprocessing was performed using a custom script that employed tools from the FMRIB Software Library (FSL) (Jenkinson et al. 2012), AFNI (Cox 1996), FreeSurfer (Fischl 2012) and MATLAB (The MathWorks Inc., Natick, Massachusetts, USA). The pCASL datasets were brain extracted (FSL bet) and split into their even and odd volumes to separate the control and tag images (FSL slicetimer and FSL fslsplit), motion correction was done separately for the even and odd images. Final BOLD maps were calculated using a surround mean difference approach (Liu and Wong 2005). CBF-fMRI data was preprocessed using ASLprep (Adebimpe et al. 2022). MATLAB was used for temporal and spatial outlier removal, drift correction and normalization. The first five volumes of both BOLD and CBF were rejected to allow the brain to enter a steady state. All datasets were registered and resampled (FSL flirt) into MNI space for group analysis.

### Statistical analysis

Following preprocessing, the data was input into FSL MELODIC (Beckmann and Smith 2004) for an independent component analysis (ICA) regardless of protocol and melanin grouping (N = 180 data sets), as shown in **Figure 2**. The ICA was chosen over the conventional general-linear model (GLM) analysis, as the hemodynamic response function for iPBM remains unknown.

**Figure 2:**
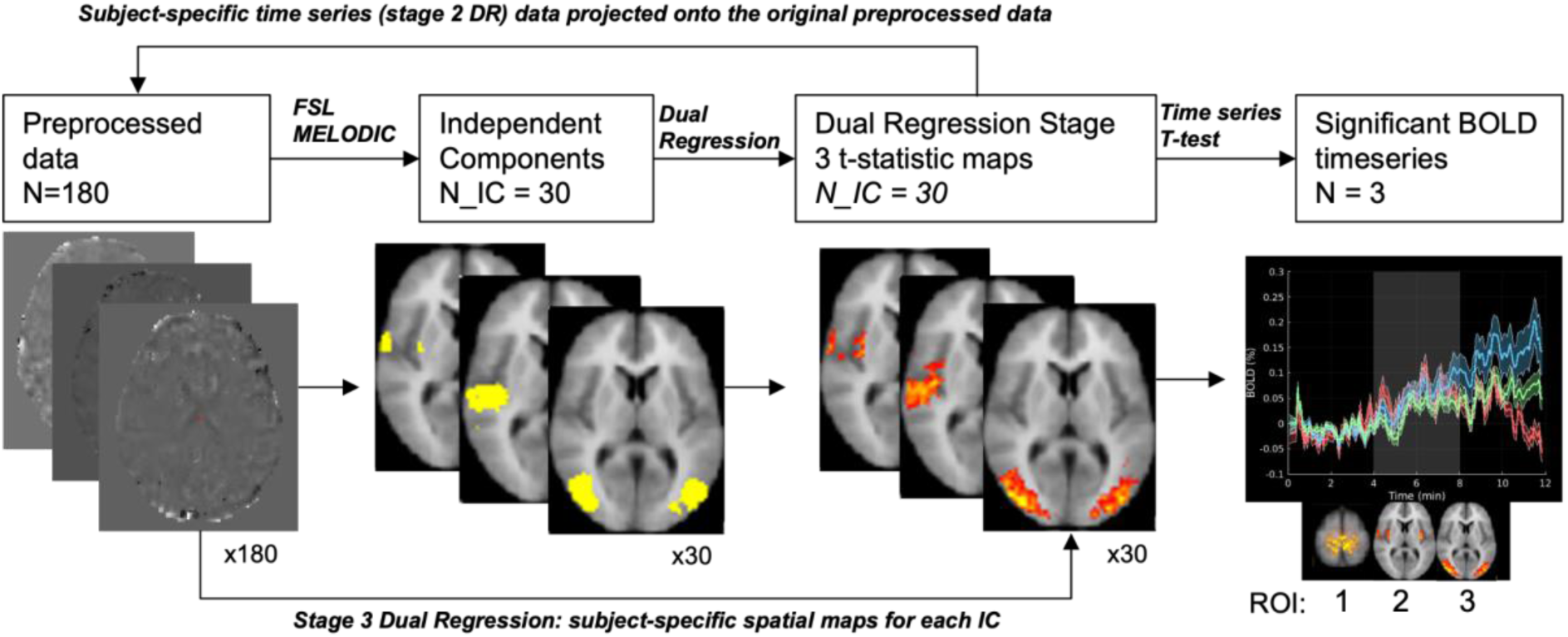
Independent component analysis and subsequent dual regression for generating the regions of interest (ROIs). All resulting independent components (ICs) were then t-tested for significance and the five remaining ICs were selected as the final ROIs.

The number of independent components (ICs) was set to 30 to avoid fragmentation. The ICs were normalized, and their temporal characteristics were examined through a t-test to determine significant correspondence with the onset of the iPBM stimulus. Subsequently, dual regression was performed to map the ICs to the normalized individual subject time series and experimental conditions. The dual regression model consists of three stages, generating statistical maps and time series for each IC, allowing for subject-specific characterization of BOLD dynamics and spatial specificity. Each of these ICs was taken as a region of interest (ROI). After dual regression, the subject-specific BOLD-normalized and CBF-normalized time series for each ROI and the site of illumination, were plotted to visualize the dynamic temporal changes in BOLD and CBF signals across conditions. To determine whether the observed BOLD and CBF signal changes were statistically significant during stimulation, we conducted a t-test on the rise of the time course within the defined experimental paradigm (4 minutes OFF, 4 minutes ON, 4 minutes OFF).

The %BOLD and %CBF change relative to the pre-stimulus period was calculated and input into a t-test for the periods during and post-stimulation. These average t-scores were incorporated into a linear mixed-effects (LME) model as the dependent variable to analyze the influence of irradiance, wavelength, frequency, melanin level (ITA) and sex on BOLD and CBF responses. However, if the time series did not exhibit a statistically significant change, the %BOLD and %CBF change for that condition was set to zero, ensuring that non-significant responses did not contribute to the statistical modeling.

The LME model followed a two-step approach. First, we used MATLAB’s stepwiselm function to identify parameter variables that had no significant effect (p >= 0.05, False Discovery Rate (FDR)-corrected) on the temporal BOLD and CBF response. These variables were removed from the final LME to maximize information content of the remaining fixed effects. The final model followed the following format, where subject was included as a random variable.

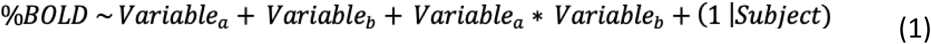

Wavelength, irradiance, pulsation frequency and sex were set as categorical variables, and ITA as a continuous variable. Final statistical p-values were thresholded through FDR correction.

### BOLD Biophysical Modeling

To further characterize the relationship between BOLD and CBF, we fit the temporal mean BOLD and CBF responses from each scan of each subject to the deoxyhemoglobin dilution model as shown in Eq. 1, with set parameters β=1.3, α=0.2. The relationship between CBF and CMRO₂ was defined by Eq. 2,

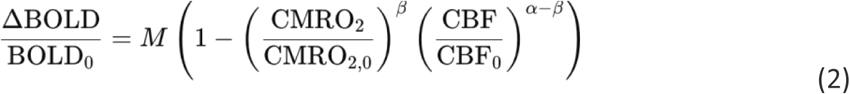

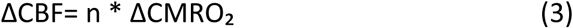

where changes in CBF and CMRO₂ are related by the neurovascular coupling ratio n. Both the n-value and M-value were derived from the above model, where the M-value estimated from the first time window was then used in the two subsequent windows to fit for the n-values. This was performed for each ROI separately

## Results

Model-free analysis across all datasets identified five regions of interests (ROIs) that corresponded to the iPBM timing vectors. These ROI spatial maps are derived from the stage 3 dual regression analysis for the ICA component associated with the iPBM timing vector. Voxels were thresholded by p < 0.05 to highlight brain regions most strongly associated with the identified component while reducing contributions from weaker or spatially diffuse effects. Further details of the ICA and dual regression methodology are provided in the supplementary materials.

Five ICs survived the ICA step and are shown in **Figure 4**, with the anatomical and functional determinants of ROIs 1-5 obtained from NeuroSynth (correlation coefficients (r) listed below) (Yarkoni et al. 2011):

- ROI 1 (**Figure 3a**) consists of subgenual (r = 0.52), ventral striatum (r=0.44) and accumbens (r = 0.40), which are associated with mood regulation, motivation and learning;
- ROI 2 (**Figure 3b**) consists of the thalamus (r = 0.28), and the caudate (r = 0.22), which are involved in sensory relay and movement execution;
- ROI 3 (**Figure 3c**) consists of the superior temporal (r = 0.63), the auditory cortex (r = 0.58), and the planum temporale (r = 0.55) which are associated with auditory and language processing;
- ROI 4 (**Figure 3d**) consists of the amygdala (r=0.28), hypothalamus (r=0.27) and the hippocampus (r=0.22), which are involved in emotional processing, regulation of bodily functions, memory and learning;
- ROI 5 (**Figure 3e**) consists of the parietal cortex (r = 0.50), superior parietal (r=0.46) and inferior parietal (r=0.43), which are associated with processing sensory information and motor control.
- ROI 6 (**Figure 3f**) represents the illuminated region, which was defined as the region most proximal to the light’s incidental site.

**Figure 3:**
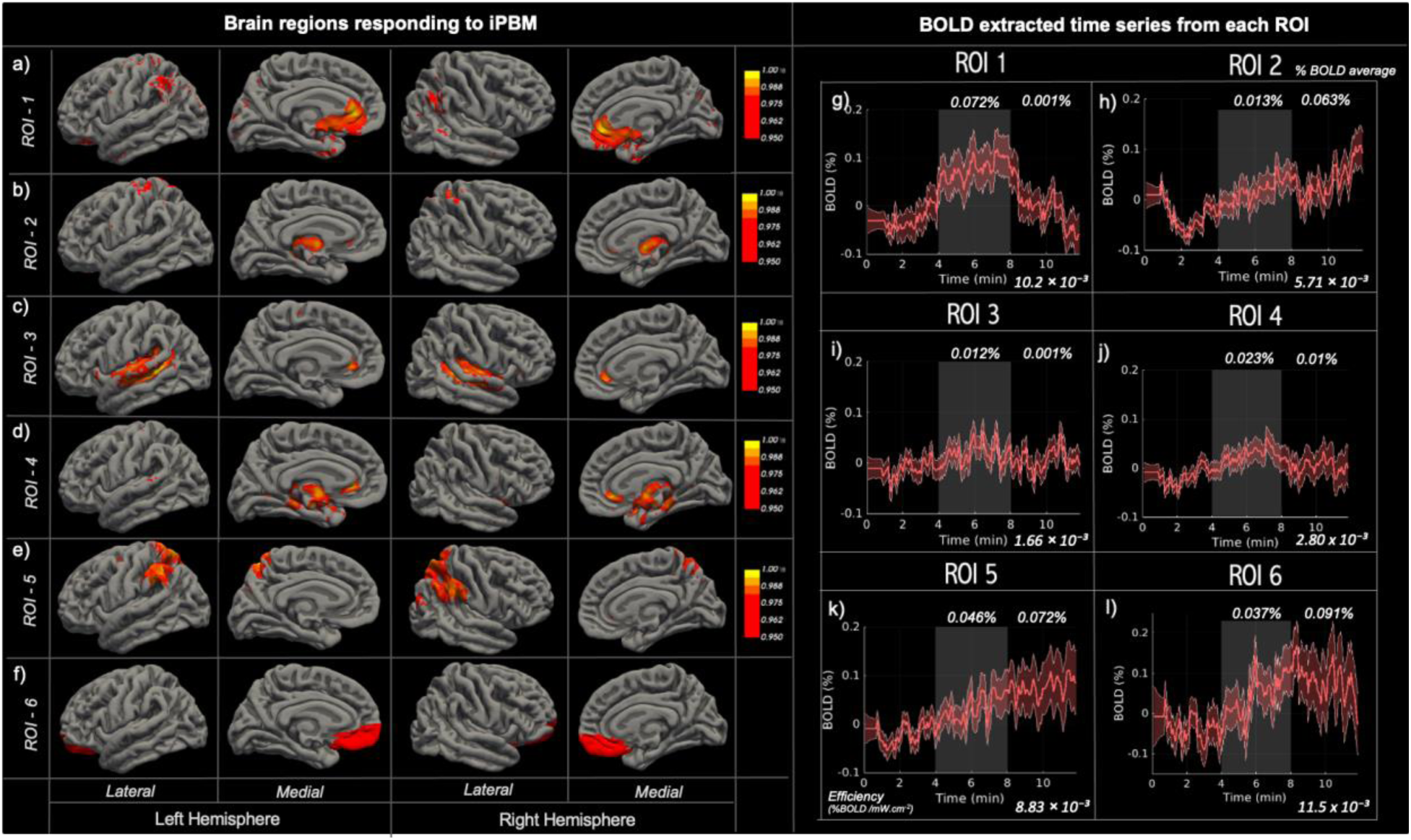
a-e) ROIs 1-5, which are Independent components showing a significant BOLD fMRI response, derived from ICA followed by dual regression, thresholded to p < 0.05. Highlighting specific brain areas most consistently associated with the PBM effects, irrespective of subject or stimulation parameter. f) ROI 6, which is the site of stimulation (illumination), manually selected based on previous Monte Carlo simulations to determine the illuminated area by iPBM, see supplemental materials section S.1. To characterize the temporal responses of each ROI, we extracted the BOLD signal trajectory in each ROI by computing the average BOLD intensity across all active voxels. Coloured shaded regions represent the standard error across all datasets. The tPBM stimulus “on” period (minutes 4:8) is denoted by the shaded grey region. The mean %BOLD increase during stimulation and post stimulation is noted for each ROI along with a metric of efficiency, determined by normalizing the largest %BOLD response by irradiance, these values are also tabulated in Table S1.

**Figure 4.**
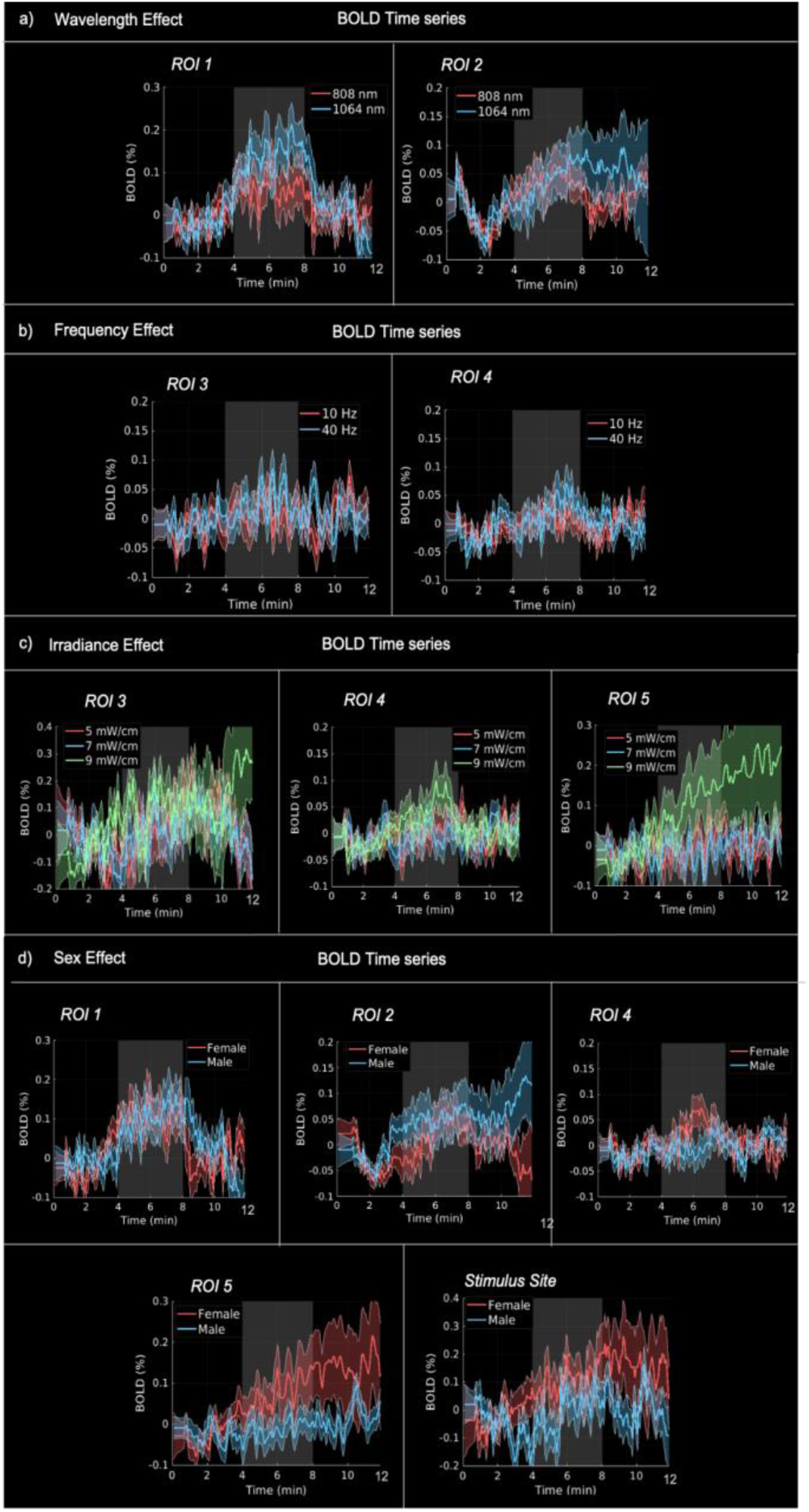
BOLD temporal response magnitudes across stimulation parameters. The y-axis represents response magnitude from the average for each ROI, reflecting the LME model response derived from 180 inputs and grouped by wavelength, irradiance, sex and frequency. All plots represent statistically significant changes (p < 0.05), with the coloured shaded regions reflecting the standard error of each timeseries. The tPBM stimulus “on” period (minutes 4:8) is denoted by the shaded grey region.

### Linear Mixed Effects Modelling

The LME modelling identified ROIs exhibiting significant differences in temporal response magnitude across various stimulation parameters; a summary of these responses is provided in **Figure 4** below.

### Biophysical Modeling

To further quantify the neurovascular mechanism underlying the observed BOLD and CBF responses using the biophysical BOLD model, we determined the slope between CBF and BOLD mean responses across participants in all six ROIs, and for different time windows in the stimulation and post-stimulation period. The individual BOLD:CBF slope for each ROI grouping is shown in **Figure 5c**. In general, regions showing increased BOLD responses also demonstrated proportional increases in CBF. This can be visualized through the scatter plots shown in **Figure 5**. Moreover, the BOLD-CBF slope declines over time, both during and after PBM stimulation.

**Figure 5:**
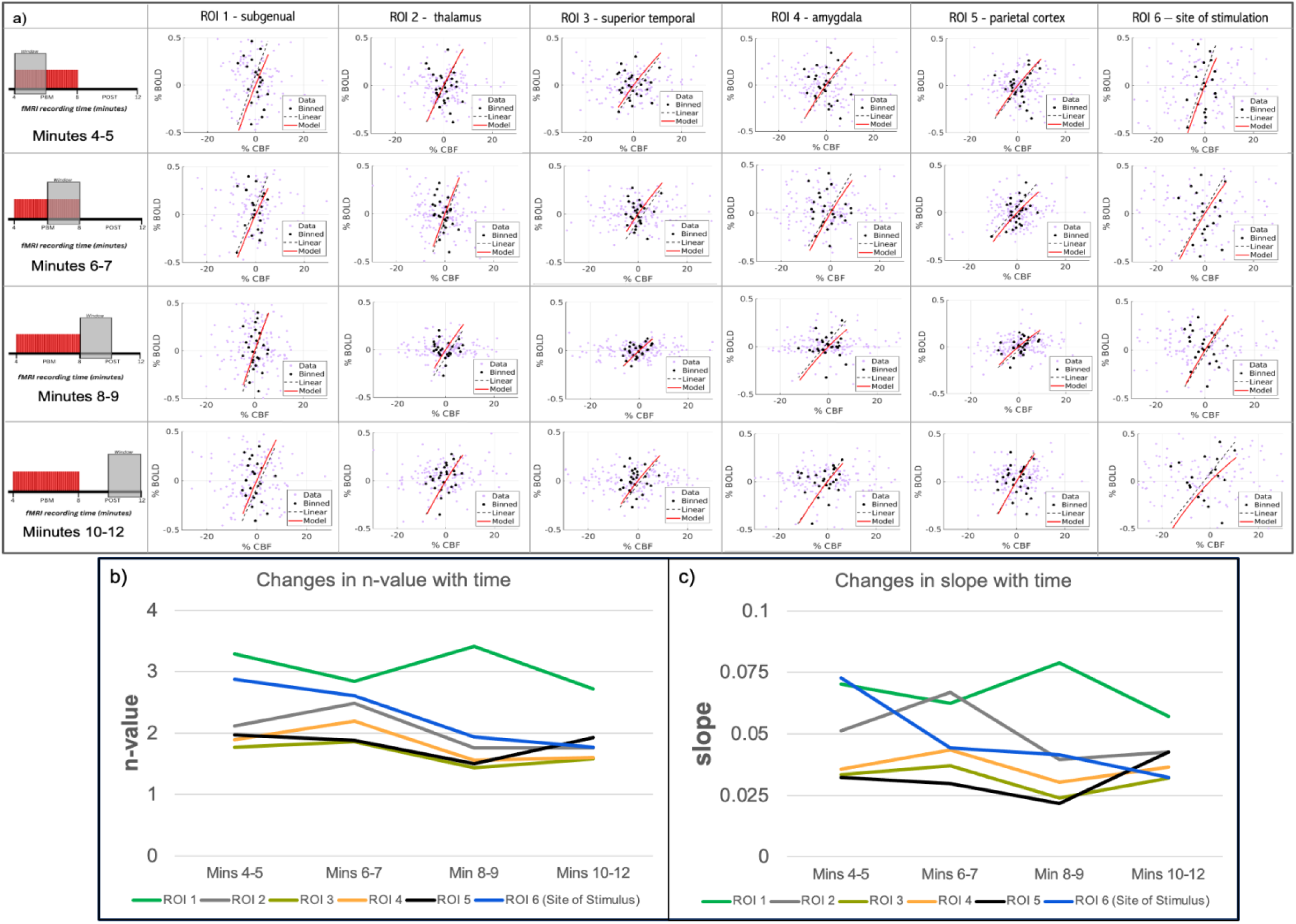
The relationship between BOLD and CBF responses across ROIs was evaluated using a neurovascular coupling model derived from the average BOLD and CBF response. (a) Scatter plot of the temporal mean BOLD and CBF responses from each scan of each subject, derived from the deoxyhemoglobin dilution model (Eq.1). The data were stratified into bins of 6 data points, and each dot represents the mean of one bin. (b) BOLD:CBF n-value variations with time for all ROIs (c) slope for each ROI, in time windows.

## Discussion

In this study, we systematically characterize the brain’s response to iPBM using human fMRI. Consistent with our hypothesis, we demonstrate the first human evidence that iPBM elicits robust and spatially structured BOLD and CBF responses, providing the grounds for comparing iPBM quantitatively to previously reported tPBM findings. Our main findings include (1) like tPBM (Van Lankveld, Chen, et al. 2025), iPBM induces a dose-dependent brain response sustained by neurovascular interactions. (2) unlike tPBM, iPBM is able to produce fMRI-detectable activity in the subcortical grey matter, (3) the iPBM brain response is highly dependent on sex, and (4) iPBM appears to exhibit much higher BOLD responses per unit of irradiance. Taken together, these findings not only establish iPBM as a viable and potentially advantageous PBM paradigm, but also highlight the potential differences between the physiological mechanisms underlying iPBM compared to tPBM.

### iPBM-induced fMRI response

Our findings demonstrate that iPBM elicits measurable BOLD fMRI responses in the brain that are comparable to those observed with more conventional paradigms such as forehead delivery (tPBM). These BOLD response amplitudes are consistent with previously reported BOLD increases during forehead tPBM (Van Lankveld, Chen, et al. 2025), albeit iPBM used a much lower range of irradiances. Moreover, the regional BOLD time courses revealed three distinct temporal profiles following iPBM; (1) stimulus-locked block-type responses, (2) ramp responses consistently rising post stimulus, and (3) ramp-type responses with quicker post-stimulus recovery (**Figure 3**). Both the block (Zhao et al. 2025) (Van Lankveld, Chen, et al. 2025) and the ramp responses (Saucedo et al. 2021; Truong et al. 2022)(Van Lankveld, Chen, et al. 2025) emulate our tPBM findings. Moreover, the subgenual cortex is a primary site of response in both iPBM and tPBM. These findings highlight that the neurovascular variations that may underlie these differences persist irrespective of the mode of PBM delivery.

The variation in the average %BOLD by ROI (**Figure 6**) suggests that the effective propagation pathway of iPBM remains undefined by simulations and may not rely solely on direct tissue penetration. iPBM stimulation elicited a response in the superior temporal cortex, which is the region furthest from the applied stimulus, that was nearly comparable in magnitude to more proximal regions. Potential mechanisms could include optical propagation through tissue, transmission via cerebrospinal fluid, or vascular-mediated distribution, highlighting the need for further investigation into the biophysical routes by which iPBM exerts its neuromodulatory effects.

**Figure 6:**
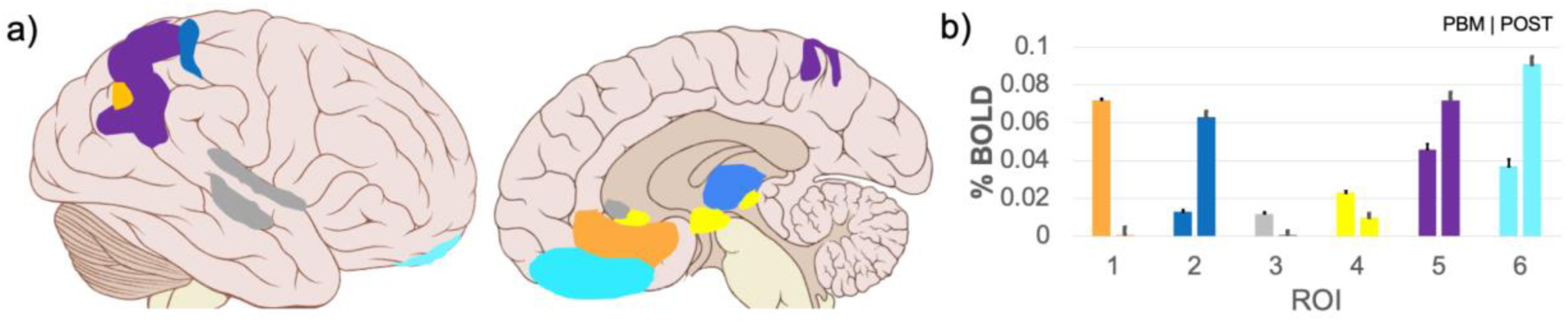
The regional responses to iPBM, and elicited %BOLD changes (a) brain rendering showing all ROIs targeted in this study, colour-coded to correspond with (b) bar graph depicting the percent BOLD change for each region, for both the during stimulus and post stimulus periods, error bars represent the standard error across all datasets.

The biophysical modeling results revealed distinct regional and temporal patterns in the coupling between BOLD and CBF responses associated with iPBM stimulation. Across regions, the slope and n-value remained positive and in consistent range with literature values, indicating that iPBM increases perfusion in excess of metabolic demand. These temporal changes show that iPBMs hemodynamic effect evolves over minutes rather than producing an immediate response. In terms of coupling dynamics, we noted that ROI 2 (thalamus) and ROI 6 (stimulus site) both exhibited a ramp-shaped response that peaked during stimulation, recovered towards baseline (at stimulus cessation) and rose again post stimulation. This response is also reflected in the BOLD-CBF coupling results (**Figure 5**), where in minutes 8-9 (immediately post stimulation) there is a drop in the n-value that then rises in the last 10-12 minutes. This multi-stage temporal profile is also consistent with the unique temporal properties of subcortical hemodynamic response functions (HRFs). While in a typical BOLD HRF, an undershoot of the signal appears after the initial peak, this feature is absent in subcortical areas (Kim et al. 2022). Supporting this, an EEG-fMRI study reported that the thalamic HRF often has two peaks compared to other cortical regions such as the parietal cortex and occipital which only had one peak (de Munck et al. 2007).

### Engaging Subcortical Regions

The significant activation in subcortical structures including the thalamus (ROI 2) and amygdala (ROI 4) we see with iPBM set it apart from tPBM. This suggests that iPBM, through the nasal cavity, may activate pathways not accessible by tPBM (due potentially to scattering and absorption throughout the scalp and skull). The cribriform plate is extremely thin compared to the frontal skull, allowing photons to penetrate with minimal energy loss. Once past this barrier, photons can directly interact with tissues that help deliver or diffuse light deeper into the brain. Photons may propagate through alternative pathways such as CSF or along the olfactory filaments and tract, effectively reaching both local and distal neural tissue targets. Subcortical regions may be activated by iPBM through both alternative pathways themselves or by triggering network activation along the pathway to the deeper structures.

The thalamus holds a central role in sensory, cognitive and motor networks, and disruption of the thalamus has been reported across multiple neurological diseases (Low et al. 2019) (Nugent et al. 2013) (Halliday 2009). The thalamus also plays a key role in recovery following traumatic brain injury (TBI) (Munivenkatappa and Agrawal 2016). On the other hand, the amygdala, a key component of the limbic system, is a primary processing center for emotions and behaviour (AbuHasan et al. 2025). Dysfunction of the amygdala is linked to anxiety, depression, PTSD and autism (Olucha-Bordonau et al. 2015).

Current non-invasive brain stimulation techniques such as transcranial magnetic stimulation (TMS) and transcranial direct current stimulation (tCDS) cannot directly stimulate deep brain structures. Recent studies have shown that stimulation of the superficial cortical regions connected to the thalamus can indirectly engage the deeper brain structures through TMS (Caparelli et al. 2022) and tDCS (Aloi et al. 2022). Emerging techniques such as transcranial focused ultrasound (TUS), can directly target deeper regions, showing thalamic stimulation enhanced connectivity in a network of sensorimotor and sensory integration areas (Kim et al. 2023). TUS has also been used to target the amygdala, revealing a decrease in amygdala activation which was correlated with decreased subjective anxiety, and overall decreased amygdala-insula, and amygdala-hippocampus resting-state connectivity <u>(Chou et al. 2024)</u>. Similar to TMS and tDCS, tPBM may be used to stimulate cortical regions that, through network connections, indirectly modulate thalamic activity. In contrast, in addition to cortical regions, iPBM offers a more effective route to deeper brain structures, directly influencing subcortical regions such as the thalamus and amygdala. This difference is relevant as it indicates that deeper brain region targets, such as the thalamus, may be reached by iPBM, offering new opportunities for clinical intervention in neurological diseases. Future studies could explore the combination of multiple stimulation strategies for best coverage and targeting.

### Dose and biological dependence of the BOLD fMRI response

The regional BOLD responses exhibited clear dose-dependent characteristics across wavelength, frequency, irradiance and sex, that were not only parameter-specific but also regionally-specific.

In terms of wavelength, 1064 nm produced larger BOLD increases than 808 nm in ROI 1 (subgenual) and ROI 2 (thalamus). Effects of pulsation frequency were also regionally-dependent: both the superior temporal cortex (ROI 3) and the amygdala (ROI 4) responded more strongly to 40 Hz compared to 10 Hz. Across ROI 4 (amygdala) and ROI 5 (parietal cortex) BOLD amplitude increased with irradiance linearly, where 9 mW/cm² produced the highest BOLD response, compared to 5 mW/cm². In ROI 3 (superior temporal cortex) an irradiance of 5 mW/cm² produced a larger BOLD response compared to 7 mW/cm².

Significant sex effects were noted in five ROIs, for ROIs 1 (subgenual), 4 (amygdala), 5 (parietal cortex) and 6 (stimulus site) females exhibited higher BOLD responses, conversely in for ROI 2 (thalamus) males showed higher BOLD responses. Sex effects were much less pronounced in tPBM (Van Lankveld, Chen, et al. 2025).

### Comparing iPBM and tPBM: Efficiency, Dose and Delivery

As alluded to earlier, we note that the iPBM mode of delivery appears to produce much stronger %BOLD changes per unit irradiance, with the efficiency being a full order of magnitude higher than tPBM (see **Table S1**). This is exemplified in the subgenual (ROI 1), which was activated in both tPBM (Van Lankveld, Chen, et al. 2025) and iPBM. This suggests that in bypassing thicker regions of the scalp and skull, the nasal pathway may allow more direct or less attenuated light delivery to ventromedial prefrontal regions, which is consistent with simulations (Cassano et al. 2019; Van Lankveld, Mai, et al. 2025). This is supported by recent light penetration simulations which have shown that iPBM can directly irradiate the ventromedial prefrontal cortex and ventromedial orbitofrontal cortex (Cassano et al. 2019; Van Lankveld, Mai, et al. 2025). Nonetheless, the simulations predicted up to 1% of the incidental energy being delivered intranasally, and based on the BOLD response, iPBM’s performance appears to go beyond confines of this theoretical prediction. This likely continues to be driven by living processes such as hemodynamics and the autonomic system (Motsenyat et al. 2025), which were discussed in detail in our previous works (Van Lankveld, Chen, et al. 2025). To conclude, the significantly lower irradiances used in iPBM eliminate the potential for thermal effects, supporting its superior safety profile for repeated and prolonged application in clinical settings.

## Limitations

This work focuses on the effect of iPBM and compares the BOLD response noted in various regions to that of our tPBM work. We note that iPBM elicits a response in subcortical structures that were not activated by tPBM, however, the specific pathways by which light reaches these regions, whether through scattering, CSF flow, blood flow or other corridors cannot be determined in this study.

Similar to our tPBM study, we employed a short 4-minute iPBM session during simultaneous BOLD and ASL fMRI, capable of detecting acute changes in physiological response but does not reveal what changes consistent light exposure over multiple days or weeks of application would induce.

Both our tPBM and iPBM work has shown consistent regions of activation that reveal parts of different known functional networks. However, the present work does not assess how the modulation of individual regions may affect the network level. As a result, while regional activations can be interpreted, the combined impact of PBM on the large-scale network dynamics remains an area for future investigation.

## Conclusion

The main findings of this study reveal that iPBM can induce measurable BOLD and CBF responses in the healthy human brain, reaching deeper structures than forehead tPBM stimulation. This measured response showed high dose dependance, with variations in optimal parameters based on stimulated region. Our work demonstrates that males generally have a lower BOLD response than females in all activated regions except the thalamus. The 1064 nm wavelength and 40 Hz pulsation frequencies dominated their respective activated regions, and irradiance was highly variable across all ROIs. In general, the iPBM response elicits a lower %BOLD than tPBM albeit at a much lower irradiance, therefore when comparing overall efficiency factoring in power density, iPBM induces a 28x greater response per unit of irradiance. This is the first study to demonstrate a robust real-time BOLD-fMRI response to iPBM comparable to that of tPBM, and in deeper structures.

## Acknowledgments

We wish to thank our funding supporters; the Ontario Centre for Innovation and the Natural Sciences and Engineering Research Council of Canada and we are grateful for financial donations from Ms. Linda Reed. Furthermore, we thank Vielight Inc. for their partnership in this work.

## Supplementary Materials

### S.1 Methods - Illumination Site

Our previous work focused on using Monte Carlo simulations of a single laser source for intranasal photobiomodulation (iPBM, nostril position) irradiation on a healthy human brain model (Colin27) (Van Lankveld, Mai, et al. 2025). For this study, an ROI of the simulated Monte Carlo region (Figure S1) was segmented as the “site of stimulation”, (ROI 6) in our analysis.

**Figure S1:**
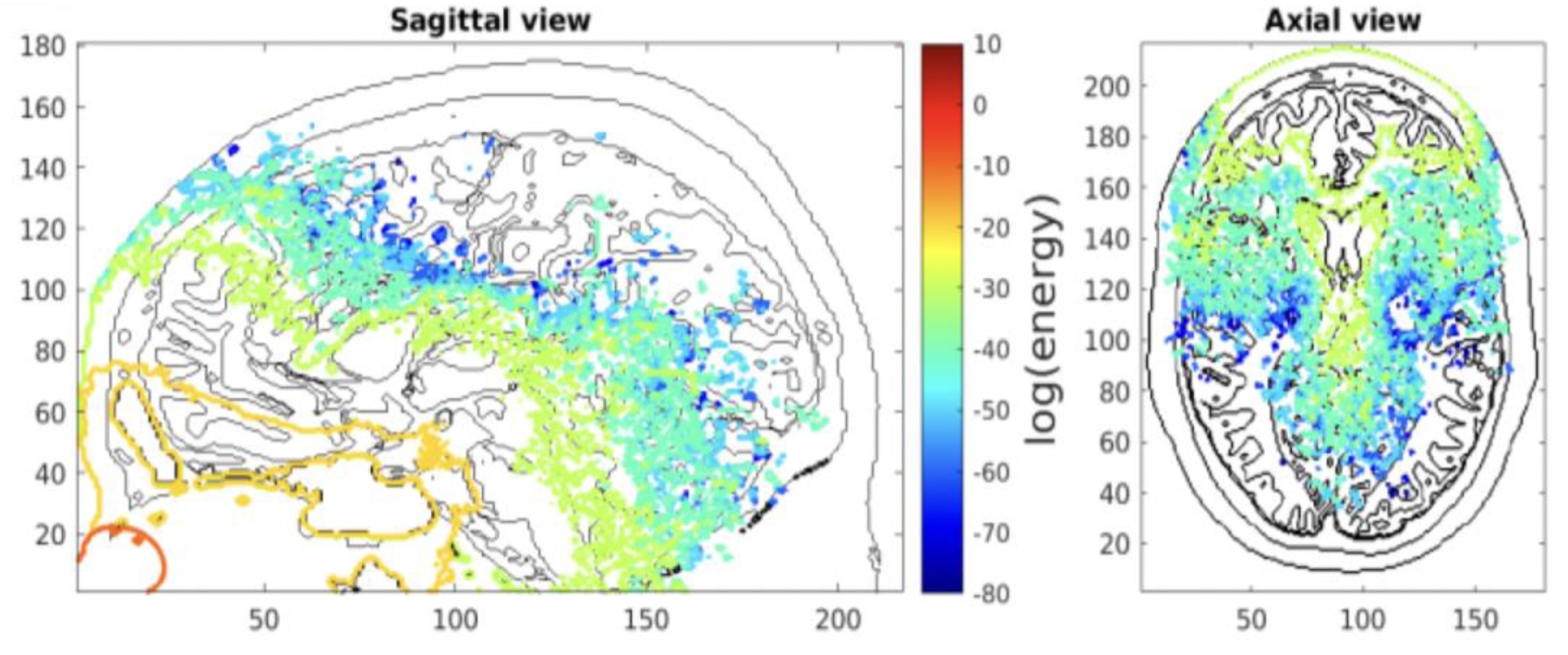
Sagittal and axial viewpoints of iPBM Monte Carlo simulated energy deposition profile, the axes represent the voxel dimensions, and the colour scale represents the log of energy levels. Figure taken from (Van Lankveld, Mai, et al. 2025).

### S.2 Results: Linear Mixed Effects Modeling Dose Dependance

**Table S1.**
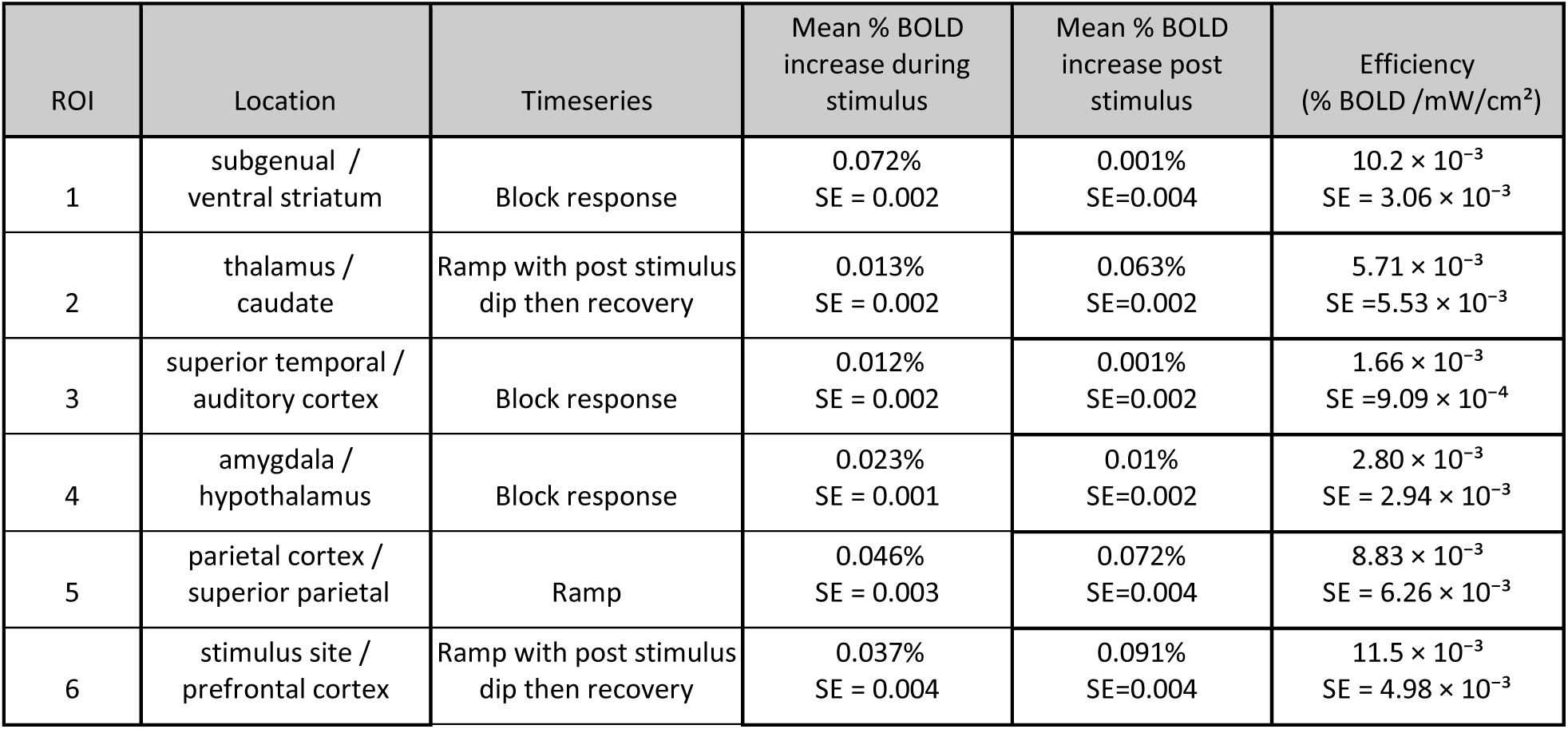
Summary of regional %BOLD measurements.

**Table S2:**
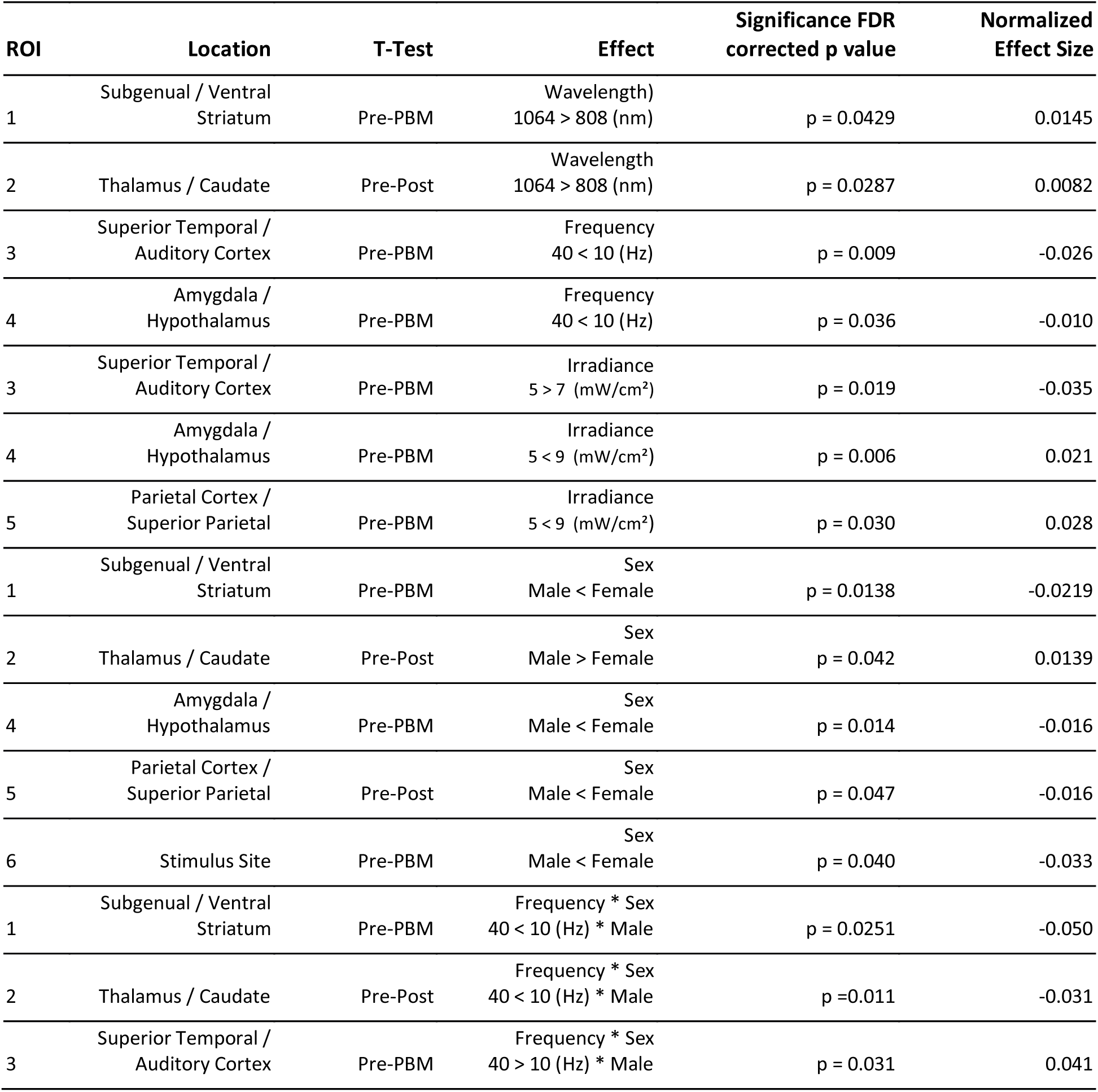
ROI-level statistical summary results from the Linear Mixed Effects model of stimulation parameters.

| ROI | Location | T-Test | Effect | Significance FDR corrected p value | Normalized Effect Size |
| --- | --- | --- | --- | --- | --- |
| 1 | Subgenual / Ventral Striatum | Pre-PBM | Wavelength<br>1064 > 808 (nm) | p = 0.0429 | 0.0145 |
| 2 | Thalamus / Caudate | Pre-Post | Wavelength<br>1064 > 808 (nm) | p = 0.0287 | 0.0082 |
| 3 | Superior Temporal / Auditory Cortex | Pre-PBM | Frequency<br>40 < 10 (Hz) | p = 0.009 | -0.026 |
| 4 | Amygdala / Hypothalamus | Pre-PBM | Frequency<br>40 < 10 (Hz) | p = 0.036 | -0.010 |
| 3 | Superior Temporal / Auditory Cortex | Pre-PBM | Irradiance<br>5 > 7 (mW/cm<sup>2</sup>) | p = 0.019 | -0.035 |
| 4 | Amygdala / Hypothalamus | Pre-PBM | Irradiance<br>5 < 9 (mW/cm<sup>2</sup>) | p = 0.006 | 0.021 |
| 5 | Parietal Cortex / Superior Parietal | Pre-PBM | Irradiance<br>5 < 9 (mW/cm<sup>2</sup>) | p = 0.030 | 0.028 |
| 1 | Subgenual / Ventral Striatum | Pre-PBM | Sex<br>Male < Female | p = 0.0138 | -0.0219 |
| 2 | Thalamus / Caudate | Pre-Post | Sex<br>Male > Female | p = 0.042 | 0.0139 |
| 4 | Amygdala / Hypothalamus | Pre-PBM | Sex<br>Male < Female | p = 0.014 | -0.016 |
| 5 | Parietal Cortex / Superior Parietal | Pre-Post | Sex<br>Male < Female | p = 0.047 | -0.016 |
| 6 | Stimulus Site | Pre-PBM | Sex<br>Male < Female | p = 0.040 | -0.033 |
| 1 | Subgenual / Ventral Striatum | Pre-PBM | Frequency * Sex<br>40 < 10 (Hz) * Male | p = 0.0251 | -0.050 |
| 2 | Thalamus / Caudate | Pre-Post | Frequency * Sex<br>40 < 10 (Hz) * Male | p = 0.011 | -0.031 |
| 3 | Superior Temporal / Auditory Cortex | Pre-PBM | Frequency * Sex<br>40 > 10 (Hz) * Male | p = 0.031 | 0.041 |

### S.3 Results: Minute by Minute CBF and BOLD responses

The BOLD data derived from the 6 ROIs are averaged across *N* = 180 datasets (4 scans/participant x 45 participants), as shown in Figure 5. CBF signal data is inherently noisier compared to BOLD-fMRI signals. We averaged the CBF signal across 1-minute time windows in order to visualize the CBF time course and reduce noise while preserving the actual temporal response. For comparison, we applied the same approach to the BOLD response, as shown in **Figure S3**.

**Figure S2:**
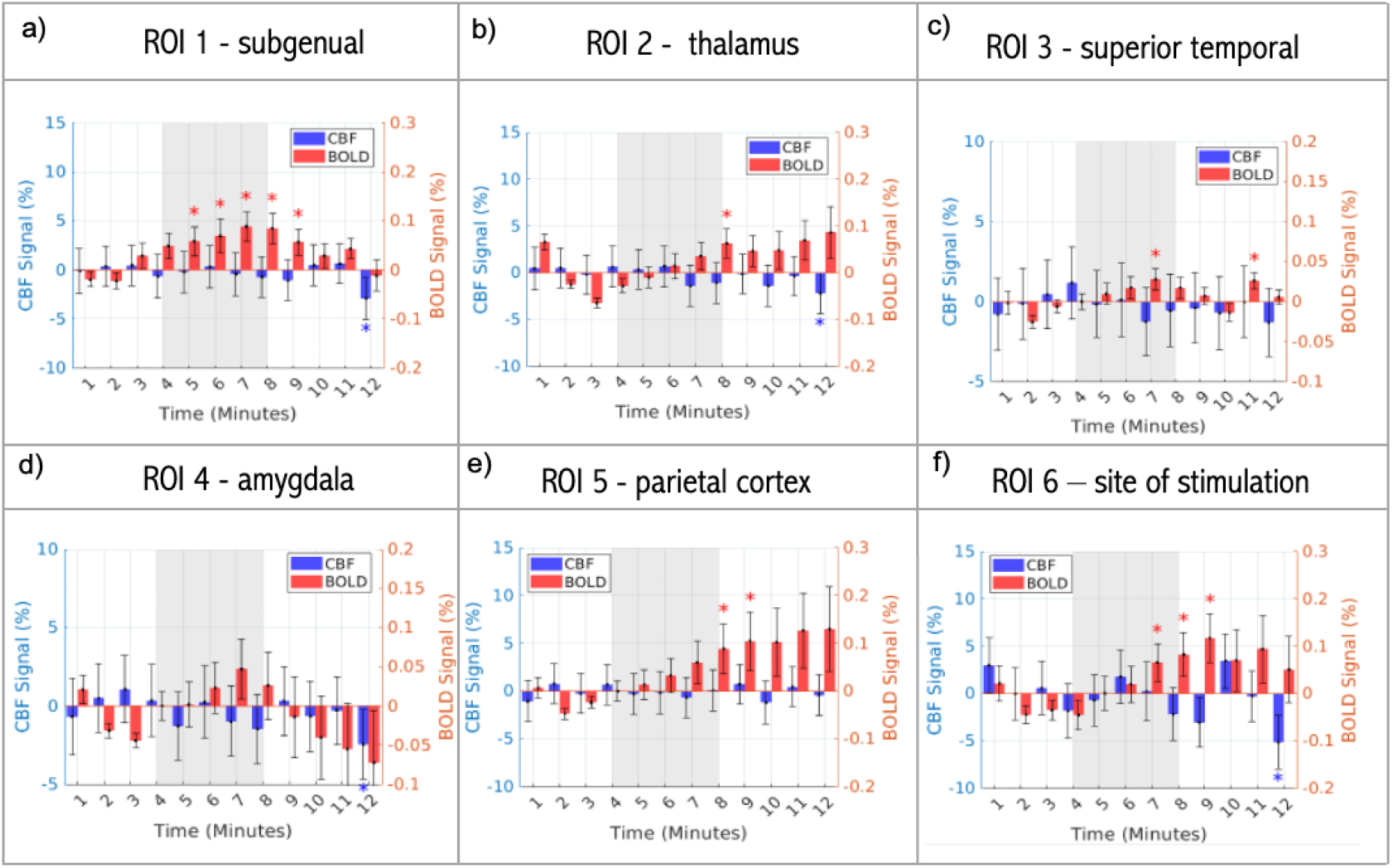
Extracted BOLD (red) and CBF (blue) time course averages for each ROI (a-e) and f) site of stimulation. Each point represents the mean signal for a 1-minute time window across all subjects and scans (n = 180) and for each ROI shown in Figure 4. Error bars represent the standard error across all datasets. The PBM stimulus (minutes 4:8) is denoted by the shaded grey region. Asterisks * indicate minutes where the response was significantly different from baseline pre-stimulus (minutes 0:4) based on a two-tailed one-sample t-test across subjects (p < 0.05)

### S.4 Results: Linear Mixed Effects Modeling

**Table S3:** BOLD response dose dependence, results from the LME. Summary of statistical results for each variable, including effect size, degrees of freedom (DoF), FDR-corrected p-value (fdr_pvalue), t-statistic (tstat), standard error (SE), and the lower and upper bounds of the confidence interval. Two contrasts were analysed the stimulus condition compared to the pre-stimulus baseline (Pre-PBM) and the post-stimulus condition compared to the pre stimulus baseline (Pre-Post)

| Pre- PBM | ROI 1 - Subgenual / Ventral Striatum |  |  |  |  |  |  |  |
| --- | --- | --- | --- | --- | --- | --- | --- | --- |
|  | Variable | Effect Size | DoF | fdr_pvalue | tstat | SE | lower bound | upper bound |
|  | Irradiance_7 | -0.002 | 156 | 0.811 | -0.240 | 0.009 | -0.019 | 0.015 |
|  | Irradiance_9 | 0.004 | 156 | 0.811 | 0.463 | 0.009 | -0.013 | 0.021 |
|  | Frequency_40 | -0.0032 | 156 | 0.7408 | -0.3314 | 0.0002 | -0.0220 | 0.0157 |
|  | Wavelength_1064 | 0.0145 | 156 | **0.0429** | 2.0411 | 0.0071 | 0.0005 | 0.0285 |
|  | Sex_Male | -0.0219 | 156 | **0.0138** | -2.4913 | 0.0088 | -0.0393 | -0.0045 |
|  | Wavelength_1064: Male | 0.0096 | 156 | 0.7510 | 0.6750 | 0.0142 | -0.0185 | 0.0377 |
|  | Frequency_40 : Male | -0.0498 | 156 | **0.0251** | -2.6719 | 0.0187 | -0.0867 | -0.0003 |
| Pre- PBM | ROI 2 - Thalamus / Caudate |  |  |  |  |  |  |  |
|  | Variable | Effect Size | DoF | fdr_pvalue | tstat | SE | lower bound | upper bound |
|  | Irradiance_7 | 0.0100 | 172 | 0.0759 | 1.7855 | 0.0056 | -0.0011 | 0.0211 |
|  | Irradiance_9 | 0.0102 | 172 | 0.0759 | 1.8343 | 0.0056 | -0.0008 | 0.0212 |
|  | Frequency_40 | 0.0001 | 172 | 0.4987 | 0.6780 | 0.0002 | -0.0002 | 0.0004 |
|  | Wavelength_1064 | 0.0066 | 172 | 0.1521 | 1.4386 | 0.0046 | -0.0024 | 0.0156 |
|  | Sex_Male | 0.0057 | 172 | 0.2181 | 1.2360 | 0.0046 | -0.0034 | 0.0147 |
|  | Wavelength_1064: Male | -0.0127 | 172 | 0.1633 | -1.4001 | 0.0090 | -0.0305 | 0.0052 |
|  | Frequency_40 : Male | -0.0071 | 172 | 0.5605 | -0.5833 | 0.0003 | -0.0312 | 0.0169 |
| Pre- PBM | ROI 3 - Superior Temporal / Auditory Cortex |  |  |  |  |  |  |  |
|  | Variable | Effect Size | DoF | fdr_pvalue | tstat | SE | lower bound | upper bound |
|  | Irradiance_7 | -0.035 | 175 | **0.019** | -2.622 | 0.013 | -0.062 | -0.009 |
|  | Irradiance_9 | 0.007 | 175 | 0.625 | 0.490 | 0.013 | -0.020 | 0.033 |
|  | Frequency_40 | -0.026 | 175 | **0.009** | -2.648 | 0.010 | -0.046 | -0.007 |
|  | Wavelength_1064 | -0.007 | 175 | 0.524 | -0.638 | 0.011 | -0.029 | 0.015 |
|  | Sex_Male | 0.016 | 175 | 0.109 | 2.473 | 0.007 | 0.003 | 0.029 |
|  | Wavelength_1064: Male | 0.003 | 175 | 0.933 | 0.171 | 0.015 | -0.027 | 0.032 |
|  | Frequency_40 : Male | 0.041 | 175 | **0.031** | 2.144 | 0.019 | 0.004 | 0.078 |
| Pre- PBM | ROI 4 - Amygdala / Hypothalamus |  |  |  |  |  |  |  |
|  | Variable | Effect Size | DoF | fdr_pvalue | tstat | SE | lower bound | upper bound |
|  | Irradiance_7 | 0.010 | 177 | 0.166 | 1.391 | 0.007 | -0.004 | 0.024 |
|  | Irradiance_9 | 0.021 | 177 | **0.006** | 2.989 | 0.007 | 0.007 | 0.034 |
|  | Frequency_40 | -0.010 | 177 | **0.036** | -2.113 | 0.005 | -0.020 | -0.001 |
|  | Wavelength_1064 | -0.003 | 177 | 0.565 | -0.577 | 0.006 | -0.015 | 0.008 |
|  | Sex_Male | -0.016 | 177 | **0.014** | -1.610 | 0.010 | -0.037 | 0.004 |
|  | Wavelength_1064: Male | -0.008 | 177 | 0.777 | -0.648 | 0.012 | -0.032 | 0.016 |
|  | Frequency_40 : Male | 0.005 | 177 | 0.843 | 0.434 | 0.012 | -0.019 | 0.030 |
| Pre- PBM | ROI 5 - Parietal Cortex / Superior Parietal |  |  |  |  |  |  |  |
|  | Variable | Effect Size | DoF | fdr_pvalue | tstat | SE | lower bound | upper bound |
|  | Irradiance_7 | -0.004 | 173 | 0.747 | -0.323 | 0.012 | -0.027 | 0.019 |
|  | Irradiance_9 | 0.028 | 173 | **0.030** | 2.459 | 0.012 | 0.006 | 0.051 |
|  | Frequency_40 | -0.015 | 173 | 0.111 | -1.601 | 0.010 | -0.034 | 0.004 |
|  | Wavelength_1064 | -0.007 | 173 | 0.494 | -0.686 | 0.010 | -0.026 | 0.012 |
|  | Sex_Male | -0.004 | 173 | 0.676 | -0.419 | 0.010 | -0.023 | 0.015 |
|  | Wavelength_1064: Male | 0.022 | 173 | 0.266 | 1.166 | 0.019 | -0.015 | 0.060 |
|  | Frequency_40 : Male | 0.015 | 173 | 0.431 | 0.789 | 0.019 | -0.023 | 0.053 |
| Pre- PBM | ROI 6 - Site of Stimulation |  |  |  |  |  |  |  |
|  | Variable | Effect Size | DoF | fdr_pvalue | tstat | SE | lower bound | upper bound |
|  | Irradiance_7 | 0.016 | 174 | 0.317 | 1.144 | 0.014 | -0.011 | 0.043 |
|  | Irradiance_9 | 0.014 | 174 | 0.317 | 1.003 | 0.014 | -0.013 | 0.040 |
|  | Frequency_40 | -0.011 | 174 | 0.343 | -0.951 | 0.012 | -0.034 | 0.012 |
|  | Wavelength_1064 | 0.007 | 174 | 0.627 | 0.486 | 0.015 | -0.023 | 0.038 |
|  | Sex_Male | -0.033 | 174 | **0.040** | -2.069 | 0.016 | -0.064 | -0.002 |
|  | Wavelength_1064: Male | 0.030 | 174 | 0.322 | 1.346 | 0.022 | -0.014 | 0.074 |
|  | Frequency_40 : Male | -0.014 | 174 | 0.830 | -0.603 | 0.023 | -0.059 | 0.031 |
| Pre- Post | ROI 1 - Subgenual / Ventral Striatum |  |  |  |  |  |  |  |
|  | Variable | Effect Size | DoF | fdr_pvalue | tstat | SE | lower bound | upper bound |
|  | Irradiance_7 | 0.0030 | 154 | 0.7973 | 0.2573 | 0.0116 | -0.0199 | 0.0259 |
|  | Irradiance_9 | 0.0197 | 154 | 0.1782 | 1.7109 | 0.0115 | -0.0031 | 0.0425 |
|  | Frequency_40 | -0.0012 | 154 | 0.9270 | -0.0918 | 0.0003 | -0.0266 | 0.0243 |
|  | Wavelength_1064 | -0.0060 | 154 | 0.5219 | -0.6419 | 0.0094 | -0.0246 | 0.0125 |
|  | Sex_Male | -0.0051 | 154 | 0.6003 | -0.5250 | 0.0097 | -0.0244 | 0.0141 |
|  | Wavelength_1064: Male | 0.0025 | 154 | 0.8965 | 0.1303 | 0.0190 | -0.0351 | 0.0401 |
|  | Frequency_40 : Male | -0.0010 | 154 | 0.9819 | -0.0402 | 0.0006 | -0.0514 | 0.0493 |
| Pre- Post | ROI 2 - Thalamus / Caudate |  |  |  |  |  |  |  |
|  | Variable | Effect Size | DoF | fdr_pvalue | tstat | SE | lower bound | upper bound |
|  | Irradiance_7 | -0.0040 | 174 | 0.7662 | -0.8035 | 0.0050 | -0.0138 | 0.0058 |
|  | Irradiance_9 | -0.0015 | 174 | 0.7662 | -0.2979 | 0.0050 | -0.0113 | 0.0083 |
|  | Frequency_40 | -0.0002 | 174 | 0.9724 | -0.0347 | 0.0002 | -0.0121 | 0.0117 |
|  | Wavelength_1064 | 0.0082 | 174 | **0.0287** | 2.2053 | 0.0037 | 0.0156 | 0.0009 |
|  | Sex_Male | 0.0139 | 174 | **0.0419** | 2.0494 | 0.0068 | 0.0005 | 0.0273 |
|  | Wavelength_1064: Male | 0.0047 | 174 | 0.7804 | 0.5932 | 0.0080 | -0.0110 | 0.0204 |
|  | Frequency_40 : Male | -0.0308 | 174 | **0.0110** | -2.9445 | 0.0003 | -0.0514 | -0.0101 |
| Pre- Post | ROI 3 - Superior Temporal / Auditory Cortex |  |  |  |  |  |  |  |
|  | Variable | Effect Size | DoF | fdr_pvalue | tstat | SE | lower bound | upper bound |
|  | Irradiance_7 | -0.024 | 176 | **0.009** | -2.870 | 0.008 | -0.040 | -0.007 |
|  | Irradiance_9 | -0.013 | 176 | 0.113 | -1.595 | 0.008 | -0.029 | 0.003 |
|  | Frequency_40 | -0.006 | 176 | 0.526 | -0.635 | 0.000 | -0.024 | 0.012 |
|  | Wavelength_1064 | 0.005 | 176 | 0.466 | 0.731 | 0.007 | -0.009 | 0.019 |
|  | Sex_Male | 0.002 | 176 | 0.741 | 0.331 | 0.007 | -0.011 | 0.016 |
|  | Wavelength_1064: Male | 0.007 | 176 | 0.919 | 0.477 | 0.014 | -0.020 | 0.034 |
|  | Frequency_40 : Male | 0.007 | 176 | 0.883 | 0.380 | 0.000 | -0.029 | 0.001 |
| Pre- Post | ROI 4 - Amygdala / Hypothalamus |  |  |  |  |  |  |  |
|  | Variable | Effect Size | DoF | fdr_pvalue | tstat | SE | lower bound | upper bound |
|  | Irradiance_7 | 0.001 | 175 | 0.937 | 0.141 | 0.007 | -0.013 | 0.016 |
|  | Irradiance_9 | 0.001 | 175 | 0.937 | 0.079 | 0.007 | -0.014 | 0.015 |
|  | Frequency_40 | 0.002 | 175 | 0.705 | 0.379 | 0.006 | -0.009 | 0.014 |
|  | Wavelength_1064 | 0.006 | 175 | 0.345 | 0.947 | 0.006 | -0.006 | 0.017 |
|  | Sex_Male | -0.003 | 175 | 0.605 | -0.518 | 0.006 | -0.015 | 0.009 |
|  | Wavelength_1064: Male | 0.021 | 175 | 0.152 | 1.806 | 0.012 | -0.002 | 0.044 |
|  | Frequency_40 : Male | 0.002 | 175 | 0.880 | 0.166 | 0.012 | -0.021 | 0.025 |
| Pre- Post | ROI 5 - Parietal Cortex / Superior Parietal |  |  |  |  |  |  |  |
|  | Variable | Effect Size | DoF | fdr_pvalue | tstat | SE | lower bound | upper bound |
|  | Irradiance_7 | 0.006 | 175 | 0.518 | 0.648 | 0.010 | -0.013 | 0.025 |
|  | Irradiance_9 | -0.025 | 175 | **0.021** | -2.593 | 0.010 | -0.045 | -0.006 |
|  | Frequency_40 | -0.014 | 175 | 0.111 | -1.602 | 0.009 | -0.031 | 0.003 |
|  | Wavelength_1064 | 0.005 | 175 | 0.636 | 0.474 | 0.010 | -0.015 | 0.024 |
|  | Sex_Male | -0.016 | 175 | **0.047** | -1.998 | 0.008 | -0.032 | 0.000 |
|  | Wavelength_1064: Male | 0.029 | 175 | 0.439 | 1.460 | 0.020 | -0.010 | 0.068 |
|  | Frequency_40 : Male | -0.003 | 175 | 0.865 | -0.170 | 0.016 | -0.035 | 0.029 |
| Pre- Post | ROI 6 - Site of Stimulation |  |  |  |  |  |  |  |

| Variable | Effect Size | DoF | fdr_pvalue | tstat | SE | lower bound | upper bound |
| --- | --- | --- | --- | --- | --- | --- | --- |
| Irradiance_7 | 0.002 | 176 | 0.919 | 0.102 | 0.022 | -0.042 | 0.047 |
| Irradiance_9 | 0.022 | 176 | 0.634 | 1.004 | 0.022 | -0.021 | 0.066 |
| Frequency_40 | 0.010 | 176 | 0.577 | 0.560 | 0.018 | -0.026 | 0.047 |
| Wavelength_1064 | 0.010 | 176 | 0.554 | 0.592 | 0.017 | -0.024 | 0.045 |
| Sex_Male | -0.051 | 176 | **0.009** | -2.631 | 0.019 | -0.090 | -0.013 |
| Wavelength_1064: Male | 0.009 | 176 | 0.836 | 0.243 | 0.036 | -0.062 | 0.080 |
| Frequency_40 : Male | 0.062 | 176 | 0.126 | 1.739 | 0.036 | -0.008 | 0.132 |

## Notes

### Competing Interest Statement

The authors have declared no competing interest.

